# Contractile work and biarticular mechanisms of the triceps surae muscles facilitate net ankle mechanical work at high walking speeds

**DOI:** 10.1101/2022.09.24.507845

**Authors:** Mohamadreza Kharazi, Sebastian Bohm, Christos Theodorakis, Falk Mersmann, Adamantios Arampatzis

## Abstract

Increasing walking speed is accompanied by an enhancement of the mechanical power and work performed at the ankle joint despite the decrease of the intrinsic muscle force potential. We measured Achilles tendon (AT) elongation and, based on an experimentally determined AT-force-elongation relationship; we quantified AT-force as a proxy of the triceps surae muscle force at four walking speeds (slow 0.7 m.s^-1^, preferred 1.4 m.s^-1^, transition 2.0 m.s^-1^ and maximum 2.6±0.3 m.s^-1^). Further, we investigated the mechanical power and work of the triceps surae muscles at the ankle joint (TSA) and the mechanical power and work of the biarticular gastrocnemii at the ankle and knee joint. We found a ~21% decrease of maximum AT-force at the two higher speeds compared to the preferred; however, the net TSA-work increased as a function of walking speed. An earlier plantarflexion accompanied by increased activation of the triceps surae muscles and a knee-to-ankle energy transfer via the biarticular gastrocnemii enhanced the net TSA-mechanical work by 1.7 and 2.4-fold in the transition and maximum walking speeds, respectively. Our findings provide first time evidence for different mechanistic participation of the monoarticular soleus muscle and the biarticular gastrocnemii for the speed-related enhancement of net TSA-work.

## Introduction

Human walking covers a wide range of speeds from ~0.4 to ~3.0 m.s^−1^ (Mademli & Arampatzis, 2014; Martin et al., 1992; Nilsson & Thorstensson, 1987). During walking, the mechanical work done at the ankle joint accounts for 40 to 50% of the total mechanical work performed in the lower extremities (D. J. Farris & G. S. Sawicki, 2012; Novacheck, 1998). Furthermore, increasing walking speed is accompanied by enhancing the mechanical power and work performed at the ankle joint (D. J. Farris & G. S. Sawicki, 2012; Lai et al., 2015). These reports show the relevant contribution of the ankle joint to the necessary mechanical power and work during walking and thus indicate the important role of the plantar flexor muscles as a key source of energy production for walking.

It is well known that the energy expenditure per meter distance or the metabolic energy cost of walking as a function of walking speed is characterized by a U-shaped curve, with a minimum at speeds around 1.4 m.s^−1^ (Browning et al., 2006; Cavagna & Kaneko, 1977; Margaria et al., 1963; Ralston, 1958; Zarrugh et al., 1974). Humans preferred walking speed is very close to the optimum speed that minimizes the metabolic energy cost of the transport (Browning et al., 2006; Martin et al., 1992). It is also accepted that humans switch voluntarily from walking to running at speeds around 2.0 m.s^−1^, although the metabolic energy cost of running is higher than that of walking (Dominic James Farris & Gregory S Sawicki, 2012; Hreljac, 1993; Minetti et al., 1994). The higher metabolic cost of running at 2.0 m.s^−1^ shows that the transition from walking to running is not primarily triggered to minimize the energy costs of locomotion. Minetti et al. (Minetti et al., 1994) suggested that the triceps surae muscles begin to work inefficiently due to the high contraction velocities at the walking-to-running transition speed. Therefore, humans switch to running to improve the efficiency of the triceps surae muscles, despite the higher metabolic energy cost. Farris and Sawicki (Dominic James Farris & Gregory S Sawicki, 2012) and Lai et al. (Lai et al., 2015) confirmed this hypothesis experimentally and found that switching from walking to running at transition speed indeed increases the efficiency of force generation of the triceps surae muscles. These results further support model predictions from Neptune and Sasaki (Neptune & Sasaki, 2005) that intrinsic muscle properties of the triceps surae might be responsible for the transition from walking to running.

Nevertheless, humans can walk at higher speeds than the transition speed, despite the above-mentioned decreased potential of the triceps surae muscles to generate forces (Mademli & Arampatzis, 2014; Nilsson & Thorstensson, 1987). The impaired force potential at walking speeds of ~2.0 m.s^−1^ (Dominic James Farris & Gregory S Sawicki, 2012; Lai et al., 2015) may suggest lower generated forces in the triceps surae muscles at higher walking speeds. Inverse dynamic approaches, however, report greater resultant ankle joint moments in walking speeds around the transition speed (2.0 to 2.1 m.s^−1^) compared to the preferred speed (de David et al., 2015; Lai et al., 2015), indicating an increased muscle force generation of the triceps surae muscles. Using the optic fiber methodology, Finni et al. (Finni et al., 1998) measured similar peak forces in the Achilles tendon (AT) at walking speeds between 1.1 and 1.8 m.s^−1^, indicating an unchanged muscle force generation of the triceps surae muscles in this range of speeds. To our knowledge, measurements of the AT forces at walking speeds higher than the transition speed have not been conducted yet. Assuming a lower triceps surae muscle force generation in the transition and higher walking speeds, and thus lower energy storage and recoil from the AT compared to the preferred speed, we can expect a modulation in the pattern of power and energy production of the triceps surae muscles for the needed mechanical power and work at the ankle joint. Using a musculoskeletal model, Neptune et al. (Neptune et al., 2008) found that the monoarticular soleus and the biarticular gastrocnemii increase the net contractile work in the transition speed compared to those that are close to the preferred one. To our knowledge, this model prediction has not yet been experimentally validated. Furthermore, there is a lack of information concerning the AT elastic energy storage and recoil and the performed musculotendinous power and work of the triceps surae muscles at maximum walking speeds.

The biarticular gastrocnemii muscles generate moments in both ankle and knee joints and thus can redistribute the generated musculotendinous power and work over the two crossing joints (Cleland, 1867; Prilutsky et al., 1996; van Ingen Schenau et al., 1987). Due to their biarticularity, the gastrocnemii muscles can also transfer power and energy from the more proximal monoarticular vastii muscles to the ankle joint (M. Bobbert et al., 1986; Gregoire et al., 1984; van Ingen Schenau et al., 1987). Biarticular muscles, therefore, may regulate the redistribution and transfer of mechanical power and energy between the crossing joints to be effective at the joint where required (Cleland, 1867; Schenau, 1989). Two separate mechanisms of the biarticular gastrocnemii muscles can affect the mechanical power and work at the ankle joint, independent of their own musculotendinous power and work production (Junius et al., 2017; Prilutsky et al., 1996; van Ingen Schenau et al., 1987). First, an energy transfer between the two joints is possible when the mechanical powers of the gastrocnemii muscles at the ankle and knee joint have opposite signs (energy transfer mechanism). Second, the gastrocnemii can simultaneously absorb or generate mechanical power and work at the two crossed joints, thus affecting the redistribution of power and work between the two joints (joint coupling mechanism). The function of the biarticular gastrocnemii muscles at the ankle and knee joints during high walking speeds and the possible modulation of biarticular mechanisms with increasing walking speeds to enhance power and work production at the ankle joint is currently not well understood.

In this study, using our recently developed methodology (Kharazi et al., 2021), we measured the AT elongation during walking (Figure 1) and, based on an experimentally determined tendon force–elongation relationship, we quantified the AT-force as a proxy of the triceps surae muscle force in four walking speeds (slow 0.7 m.s^−1^, preferred 1.4 m.s^−1^, transition 2.0 m.s^−1^ and maximum 2.6±0.3 m.s^−1^). Further, we measured the electromyographic activity (EMG) of the soleus (Sol), gastrocnemius medialis (GM) and tibialis anterior (TA) muscles, and the AT elastic strain energy storage and recoil were calculated. Finally, we investigated the mechanical power and work of the triceps surae muscles at the ankle joint and the mechanical power and work of the biarticular gastrocnemii muscles at the ankle and knee joints. Our purpose was to gain a better understanding of the monoarticular and biarticular mechanisms within the triceps surae muscles that contribute to the increased production of mechanical power and work at the ankle joint from slow to maximum walking speeds. We expected an increased net mechanical work done at the ankle joint from the triceps surae muscles as a function of walking speed, and we hypothesized a different mechanistic contribution of the monoarticular soleus and the biarticular gastrocnemii muscles to the increased net ankle joint mechanical work. More precisely, we predicted a speed-related increase in contractile net work production of the soleus muscle and an increased contribution of biarticular mechanisms for the gastrocnemii muscles from slow to maximum walking speeds.

**Figure 1.**
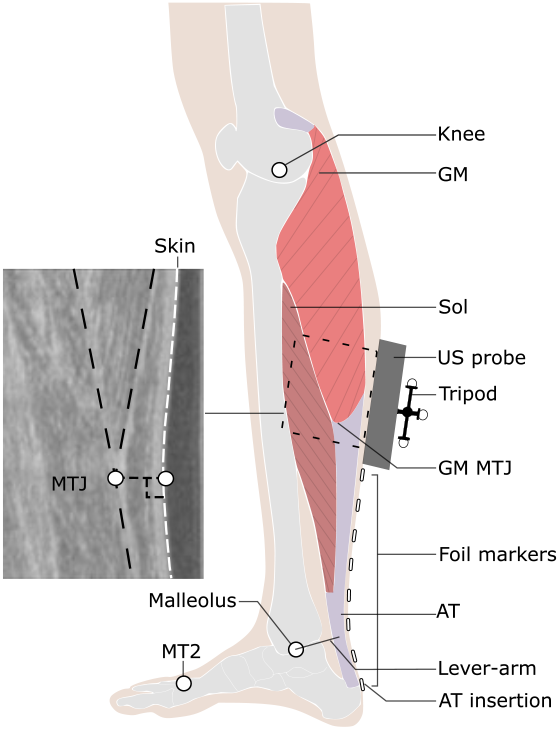
Experimental setup for determination of Achilles tendon (AT) length during locomotion. Reflective foil markers were used to reconstruct the curved shape of the AT. An ultrasound (US) probe was used to detect the movements of gastrocnemius medialis myotendinous junction (GM-MTJ), which was then projected to the skin surface (white dashed line). The tripod markers were used to transfer the detected positions of the GM-MTJ to the global coordinate system (same as the motion capture system). The AT-length was calculated as the sum of Euclidian distances from AT-insertion (notch of calcaneus bone) between every two consecutive foil markers until the projected position of GM-MTJ to the skin surface. The AT lever-arm was defined as the perpendicular distance between the midline of AT curved path and the ankle joint center. MT2: the tip of the second metatarsal; Sol: soleus muscle; Knee: a reflective marker on epicondyle of the femur.

## Results

We found a significant main effect of walking speed on all temporal and spatial gait parameters (Table 1, p<0.001). Both stance and swing time was reduced significantly (p<0.001) at higher walking speed. Step length and cadence increased as a function of speed (p<0.001), and the duty-factor decreased significantly (p<0.001) with increasing speed, however, it remained above 0.5 (i.e., the threshold for walking) at all walking speeds (Table 1). Although the time of the dorsiflexion phase decreased significantly (p<0.001) as a function of speed, the plantar flexion time remained quite preserved (Table 1).

**Table 1.** Spatial and temporal gait parameters at all walking speeds. All values are presented as mean ± standard error (fifteen individuals and nine complete gait cycles). * Statistically significant effect of speed (p<0.05). Each row sharing the same letter does not differ significantly (p>0.05, post-hoc-analysis).

| | Slow (0.7 m.s<sup>-1</sup>) | Preferred (1.4 m.s<sup>-1</sup>) | Transition (2 m.s<sup>-1</sup>) | Maximum (2.6 ± 0.3 m.s<sup>-1</sup>) |
| --- | --- | --- | --- | --- |
| Stance time (ms) * | 863±27 **a** | 601±11 **b** | 477±09 **c** | 363±11 **d** |
| Dorsiflexion time (ms) | 652±15 **a** | 416±20 **b** | 229±13 **c** | 167±06 **d** |
| Plantar flexion time (ms) | 211±14 **ab** | 185±15 **a** | 249±19 **b** | 197±09 **a** |
| Swing time (ms) * | 490±11 **a** | 411±08 **b** | 355±06 **c** | 316±5 **d** |
| Step length (m) * | 0.47±0.01 **a** | 0.7±0.01 **b** | 0.83±0.01 **c** | 0.89±0.02 **d** |
| Cadence (steps/s) * | 1.49±0.03 **a** | 1.98±0.03 **b** | 2.41±0.04 **c** | 2.95±0.06 **d** |
| Duty-factor * | 0.64±0 **a** | 0.59±0 **b** | 0.57±0 **c** | 0.53±0.01 **d** |

Ankle and knee joint angles (Figure 2B and C), strain (Figure 2A), force (Figure 2D) and lever-arm (Figure 2E) of the AT and ankle joint moments generated by the triceps surae muscles (TSA-moment, Figure 2F) during the stance phase of all walking speeds are shown in Figure 2. There was a significant main effect of walking speed on maximum AT-strain (p=0.014; Table 2) and AT-force (p=0.005; Table 2). The post hoc comparisons showed a lower maximum AT-force in the transition (p=0.004) and maximum walking speed (p=0.047) compared to the preferred one. The maximum AT-strain was lower in the transition compared to the preferred speed (p=0.017, Table 2) and showed a tendency (p=0.058) for lower values in the maximum walking speed compared to the preferred one. We found a main effect of speed on the average AT lever-arm (p=0.003).

**Figure 2.**
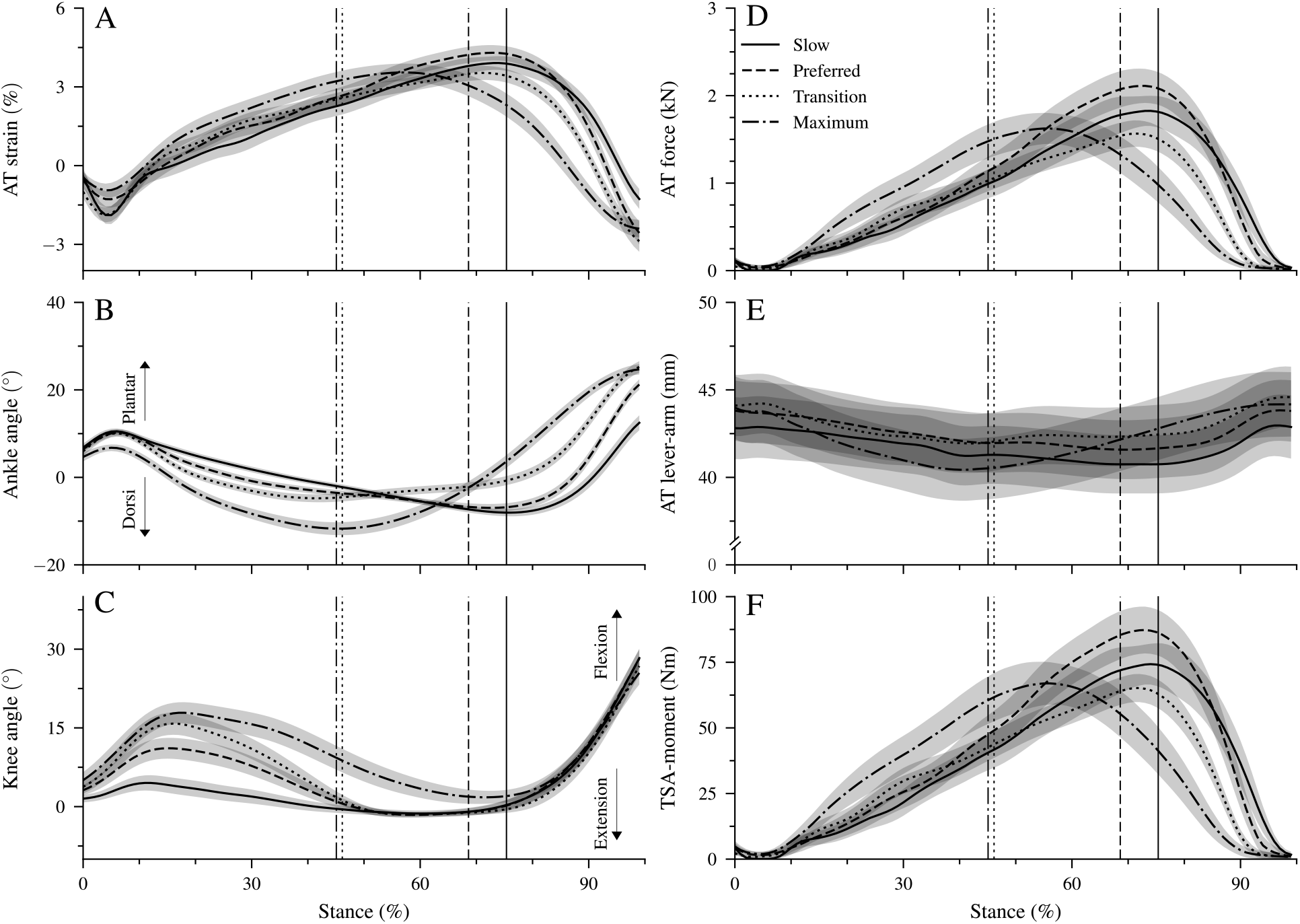
Achilles tendon (AT) strain (A), ankle and knee joint angles (B and C), AT force (D), AT lever-arm (E) and moment generated from the triceps surae muscles at the ankle joint (TSA-moment, F) during the stance phase of walking at slow (0.7 m.s^−1^, Slow), preferred (1.4 m.s^−1^, Preferred), transition (2 m.s^−1^, transition) and maximum speed (2.6±0.3 m.s^−1^, Max). The vertical solid, dashed, dotted, and dashed-dotted lines separate the dorsiflexion and plantar flexion of the ankle during walking at slow, preferred, transition and maximum speed, respectively. The curves and shaded areas represent mean ± standard errors (average of fifteen participants with nine gait cycles).

**Table 2.** Peak values of the Achilles tendon (AT) strain and force, average AT lever-arm and maximum moment generated by the triceps surae muscles at the ankle joint (TSA-moment) at all walking speeds. All values are presented as mean ± standard error (fifteen individuals and nine complete gait cycles). * Statistically significant effect of speed (p<0.05). Each row that shares the same letter does not differ significantly (p>0.05, post-hoc-analysis).

| | Slow (0.7 m.s<sup>-1</sup>) | Preferred (1.4 m.s<sup>-1</sup>) | Transition (2 m.s<sup>-1</sup>) | Maximum (2.6 ± 0.3 m.s<sup>-1</sup>) |
| --- | --- | --- | --- | --- |
| AT-strain (%) * | 4.1±0.3 **ab** | 4.4±0.3 **a** | 3.6±0.3 **b** | 3.8±0.4 **ab** |
| AT-force (N) * | 1930±190 **ab** | 2174±202 **a** | 1616±147 **b** | 1793±2.3 **b** |
| AT lever-arm (mm) * | 41±1 **a** | 42±1 **ab** | 43±1 **b** | 42±1 **a** |
| TSA-moment (Nm) * | 78±08 **ab** | 90±09 **a** | 67±05 **b** | 74±09 **ab** |

The AT lever-arm in the transition speed was longer than slow (p=0.002) and maximum (p=0.020) walking speeds (Figure 2E). However, the differences were rather small with ~1 mm (i.e., 2% of AT lever-arm, Table 2). There was a significant main effect of walking speed (p<0.001) on the maximum TSA-moment (Table 2).

Post hoc comparisons demonstrated a lower maximum TSA-moment in the transition compared to the preferred (p=0.005, Table 2) speed and a tendency for lower values in maximum walking (p = 0.050). In the slow and preferred speed, the initiation of the ankle plantar flexion and knee flexion in the second part of the stance phase occurred almost simultaneously (Figure 2B and C). In the two higher walking speeds, the time-wise earlier initiation of the plantar flexion was combined with a knee extension. During the simultaneous plantar flexion and knee extension, the knee joint extended 7.5±1.2° in the transition and 8.7±1.3° in the maximum walking speeds.

The mechanical power (Figure 3A-D) and work (Figure 3E-H) performed at the ankle joint from the triceps surae muscles (TSA-power/work) and EMG-activity of Sol, GM and TA muscles (Figure 3I-L) are presented in Figure 3. Speed affected the TSA-work during both dorsiflexion and plantar flexion periods (p<0.001, Table 3). The TSA-work during the dorsiflexion was the lowest in the transition speed, and during the plantar flexion, the highest in the maximum speed (p<0.001, Figure 3F, H and Table 3). The net TSA-work demonstrated a significant main effect of speed (p<0.001) with a continuous increase as a function of walking speed (Figure 3E-H and Table 3). The maximum AT strain energy storage and recoil was also affected by walking speed (p<0.05).

**Figure 3.**
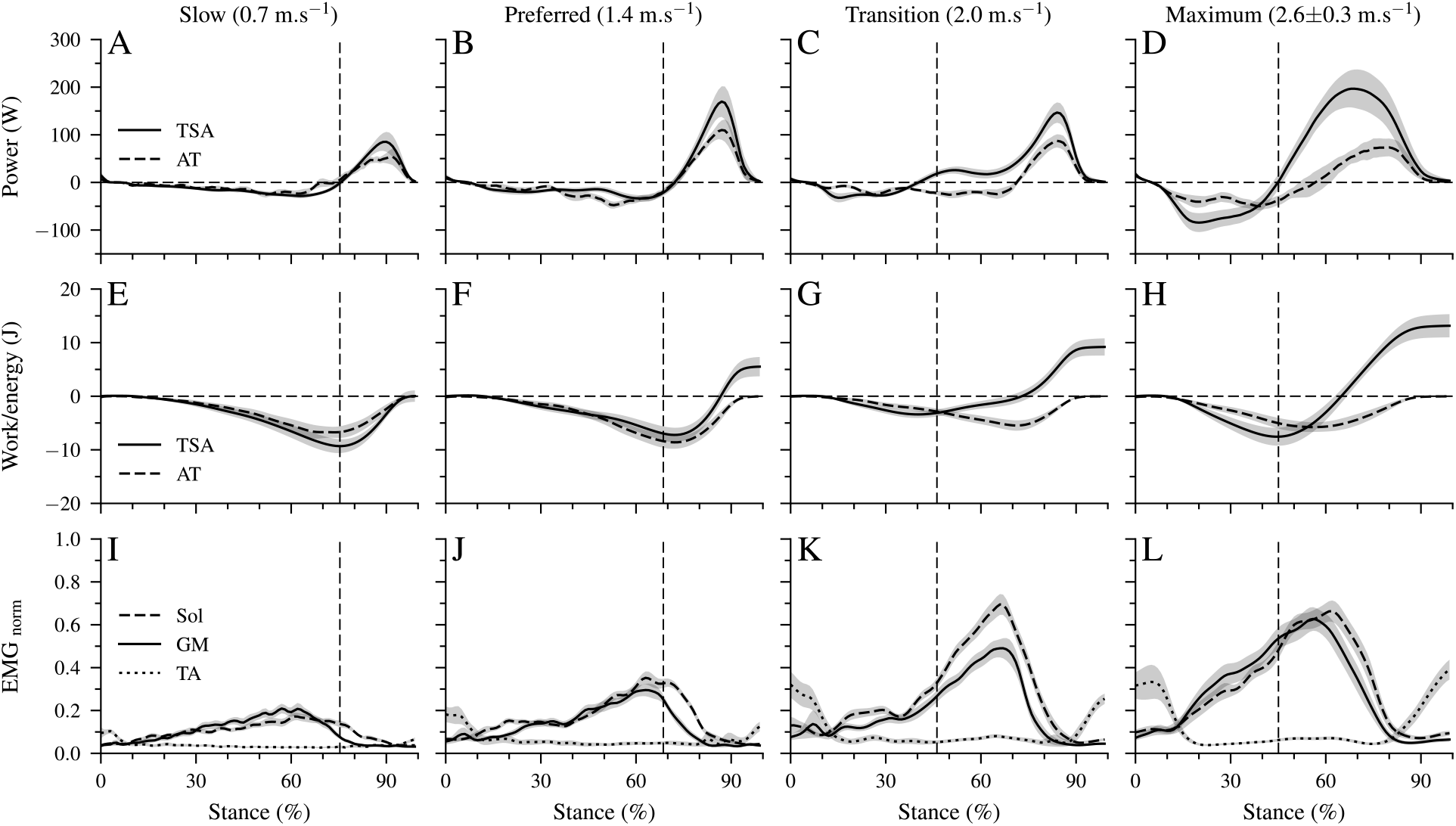
Mechanical power (A-D) and work (E-H) performed at the ankle joint by the triceps-surae muscles (TSA-power/work) as well as Achilles tendon power (AT-power) and elastic strain energy storage and recoil (AT-energy). Negative values in the mechanical power indicate energy absorption at the ankle joint during dorsiflexion and AT elastic strain energy storage. Positive power values indicate energy production at the ankle joint during the plantarflexion and elastic AT strain energy recoil. Panel I to L show the electromyographic activity of the soleus (Sol), gastrocnemius medialis (GM) and tibialis anterior (TA) muscles normalized to a maximum voluntary contraction (EMG_norm_). All investigated parameters are presented for the stance phase of walking at slow (0.7 m.s^−1^), preferred (1.4 m.s^−1^), transition (2 m.s^−1^) and maximum speed (2.6±0.3 m.s^−1^) as mean ± standard error (average of fifteen participants with nine gait cycles). The vertical dashed line shows the separation between dorsiflexion and plantar flexion.

**Table 3.** Mechanical work of triceps surae muscles on the ankle joint (TSA) during dorsiflexion, plantar flexion, and net TSA mechanical work. Achilles tendon energy storage and recoil (AT energy) and normalized maximum electromyographic activity of the soleus (Sol), gastrocnemius medialis (GM) and tibialis anterior (TA). All values are presented as mean ± standard error (fifteen individuals and nine complete gait cycles). * Statistically significant effect of speed (p<0.05). Each row that shares the same letter does not differ significantly (p>0.05, post-hoc-analysis).

| | Slow ($0.7 \text{ m.s}^{-1}$) | Preferred ($1.4 \text{ m.s}^{-1}$) | Transition ($2 \text{ m.s}^{-1}$) | Maximum ($2.6 \pm 0.3 \text{ m.s}^{-1}$) |
| --- | --- | --- | --- | --- |
| TSA dorsiflexion (J) * | $-9.6 \pm 1.3$ **a** | $-8.0 \pm 1.2$ **a** | $-4.0 \pm 0.7$ **b** | $-8.1 \pm 1.7$ **a** |
| TSA plantarflexion (J) * | $9.6 \pm 2.0$ **a** | $13.5 \pm 2.5$ **a** | $13.17 \pm 2.0$ **a** | $21.3 \pm 3.2$ **b** |
| TSA net (J) * | $0.0 \pm 1.1$ **a** | $5.5 \pm 1.8$ **b** | $9.19 \pm 1.6$ **c** | $13.2 \pm 2.1$ **d** |
| AT energy (stored/recoiled, J) * | $7.2 \pm 1.1$ **ab** | $8.8 \pm 1.1$ **a** | $5.66 \pm 0.9$ **b** | $6.7 \pm 1.1$ **ab** |
| Sol * | $0.20 \pm 0.02$ **a** | $0.39 \pm 0.02$ **b** | $0.73 \pm 0.04$ **c** | $0.77 \pm 0.04$ **c** |
| GM * | $0.25 \pm 0.02$ **a** | $0.35 \pm 0.03$ **a** | $0.61 \pm 0.06$ **b** | $0.71 \pm 0.06$ **c** |
| TA * | $0.13 \pm 0.02$ **a** | $0.22 \pm 0.04$ **a** | $0.38 \pm 0.06$ **b** | $0.51 \pm 0.06$ **c** |

The energy storage and recoil were significantly lower in the transition (p=0.011) and showed a tendency for lower values in the maximum speed (p=0.084) compared to the preferred one (Table 3). The AT strain energy recoil was significantly higher than the net TSA-work in the slow (p<0.001) and preferred (p=0.01) speed, while in the two higher speeds, the net TSA-work was 1.6 to 1.9-fold greater than the AT energy recoil. The maximum EMG-activity of all three investigated muscles demonstrated a significant speed effect (p<0.001) with the highest values in the transition and maximum speeds (p<0.001; Table 3). Figure 4 shows the mechanical power and work performed at the ankle and knee joint by the biarticular gastrocnemii muscles and the total mechanical power and work of the gastrocnemii muscle-tendon unit (MTU). The absorbed and generated work at the ankle and knee joint by the gastrocnemii muscles showed a significant speed effect (p<0.001, Table 4). The absorbed work at the ankle joint was the lowest in the transition speed, and the generated work was the highest in the maximum walking speed (p<0.001, Table 4).

**Figure 4.**
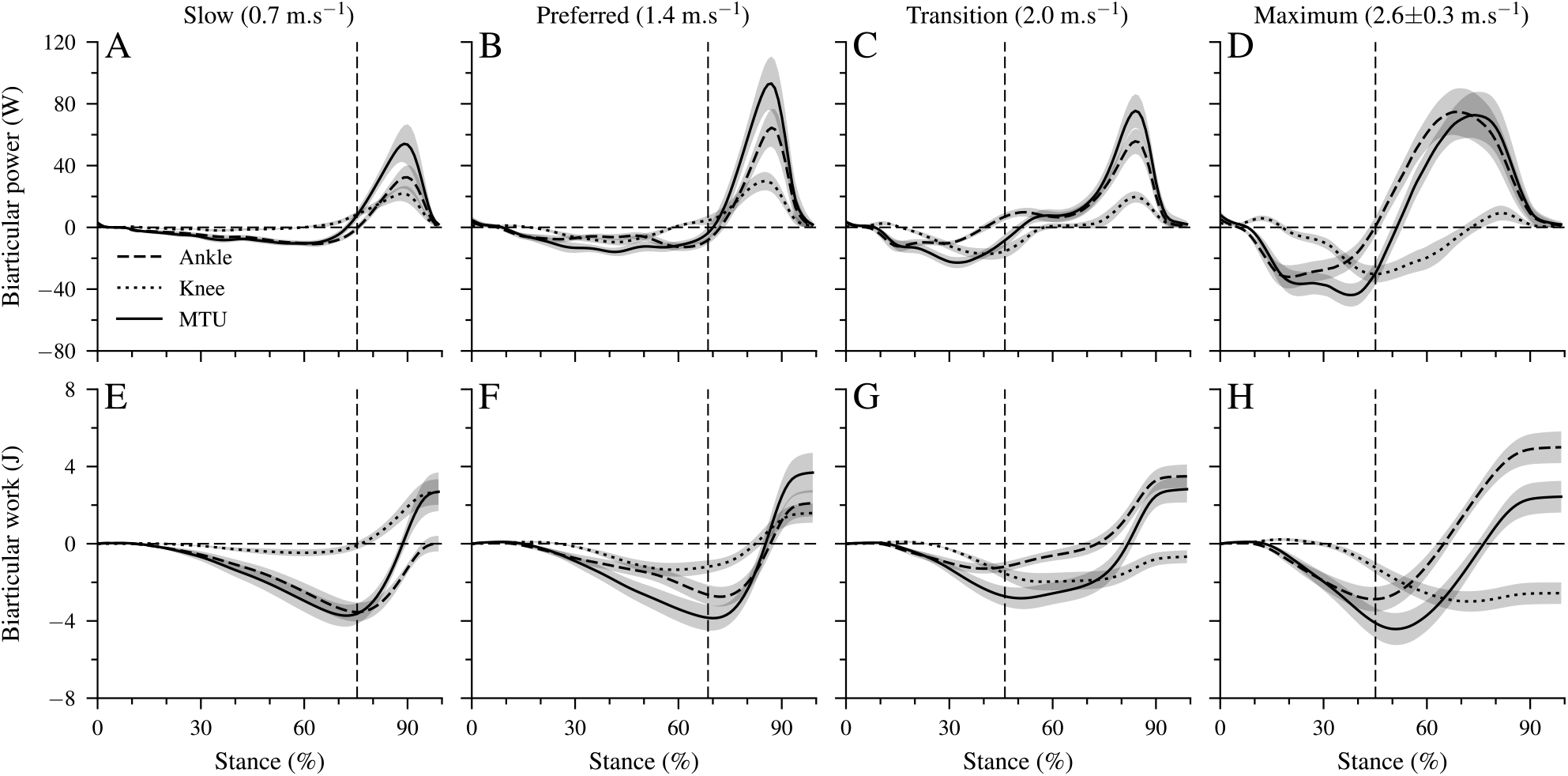
Mechanical power (A-D) and work (E-H) of the biarticular gastrocnemii muscles on the ankle and knee joints, as well as the total mechanical power/work of the gastrocnemii muscle-tendon unit (MTU). Negative values in the mechanical power indicate energy absorption at the ankle joint during dorsiflexion, energy absorption at the knee joint during knee extension and energy absorption of the MTU during lengthening. Positive power values indicate energy generation during ankle plantar flexion, knee flexion and MTU shortening. All investigated parameters are presented for the stance phase of walking at slow (0.7 m.s^−1^), preferred (1.4 m.s^−1^), transition (2 m.s^−1^) and maximum speed (2.6±0.3 m.s^−1^) as mean ± standard error. The vertical dashed line shows the separation between dorsiflexion and plantar flexion.

**Table 4.** The absorbed, generated and net work of the gastrocnemii muscles at the ankle and knee joints and the gastrocnemii muscle-tendon unit (MTU). All values are presented as mean ± standard error (fifteen individuals and nine complete gait cycles). * Statistically significant effect of speed (p<0.05). Each row sharing the same letter does not differ significantly (p>0.05, post-hoc-analysis)

| | Gastrocnemii muscles | Slow (0.7 m.s<sup>-1</sup>) | Preferred (1.4 m.s<sup>-1</sup>) | Transition (2 m.s<sup>-1</sup>) | Maximum (2.6 ± 0.3 m.s<sup>-1</sup>) |
| --- | --- | --- | --- | --- | --- |
| Ankle | Absorbed (J) * | -3.7±0.5 **a** | -3.1±0.5 **a** | -1.5±0.3 **b** | -3.1±0.7 **a** |
| | Generated (J) * | 3.8±0.8 **a** | 5.2±1.0 **a** | 5.0±0.8 **a** | 8.1±1.2 **b** |
| | Net (J) * | 0.1±0.4 **b** | 2.1±0.7 **a** | 3.5±0.6 **c** | 5.0±0.8 **d** |
| Knee | Absorbed (J) * | -0.6±0.1 **b** | -1.7±0.4 **a** | -2.5±0.5 **a** | -3.4±0.6 **c** |
| | Generated (J) * | 3.4±0.7 **a** | 3.3±0.6 **a** | 1.8±0.3 **b** | 0.9±0.3 **c** |
| | Net (J) * | 2.7±0.7 **a** | 1.6±0.5 **a** | -0.6±0.3 **b** | -2.5±0.5 **c** |
| MTU | Absorbed (J) | -3.9±0.6 | -4.2±0.7 | -3.0±0.6 | -4.6±0.9 |
| | Generated (J) | 6.6±1.3 | 7.9±1.4 | 5.9±0.9 | 7.1±1.1 |
| | Net (J) | 2.7±1.0 | 3.7±1.0 | 2.8±0.7 | 2.4±0.8 |

The net mechanical work performed at the ankle joint from gastrocnemii muscles increased significantly (p<0.001) from slow to maximum walking speed (Table 4). At the knee joint, the absorbed work increased, and the generated work decreased by increasing walking speed (p<0.05, table 4). As a result, the net mechanical work of the gastrocnemii at the knee joint decreased as a function of speed (p<0.001). The absorbed (p=0.098), generated (p=0.149), and net (p=0.323) mechanical work of the gastrocnemii MTU did not show a significant speed effect (Table 4).

There were four characteristic phases during the stance phase of walking where the unique property of the biarticular gastrocnemii muscles to act simultaneously on the ankle and knee joint influenced the net mechanical work at the ankle joint despite the unchanged net mechanical work of their MTU (Figure 2B, C and Figure 4A-D). First, at the beginning of the stance phase and during the knee flexion, power and energy were transferred from the dorsiflexed ankle joint to the knee joint (phase T1, energy transfer mechanism). Second, in the phase where the ankle joint continued the dorsiflexion and the knee joint was extended, the gastrocnemii absorbed energy in both the ankle and knee joint (phase C1, joint coupling mechanism). Third, during a simultaneous plantar flexion and knee extension, particularly in the transition and maximum walking speeds, energy was transferred from the knee to the ankle joint (phase T2, energy transfer mechanism). Fourth, at the end of the stance phase and during the synchronous plantarflexion and knee flexion, the gastrocnemii generated work in both the ankle and knee joint (phase C2, joint coupling mechanism). Figure 5A visualize the four phases, showing the mechanical power generated by the biarticular gastrocnemii muscles at the knee joint during maximum walking speed. The mechanical knee joint work in the four phases and the net knee work during all walking speeds is shown in Figure 5B. The net mechanical work performed at the knee joint determines the difference in the net work between the gastrocnemii MTU and the ankle joint and is a result of the biarticularity.

**Figure 5.**
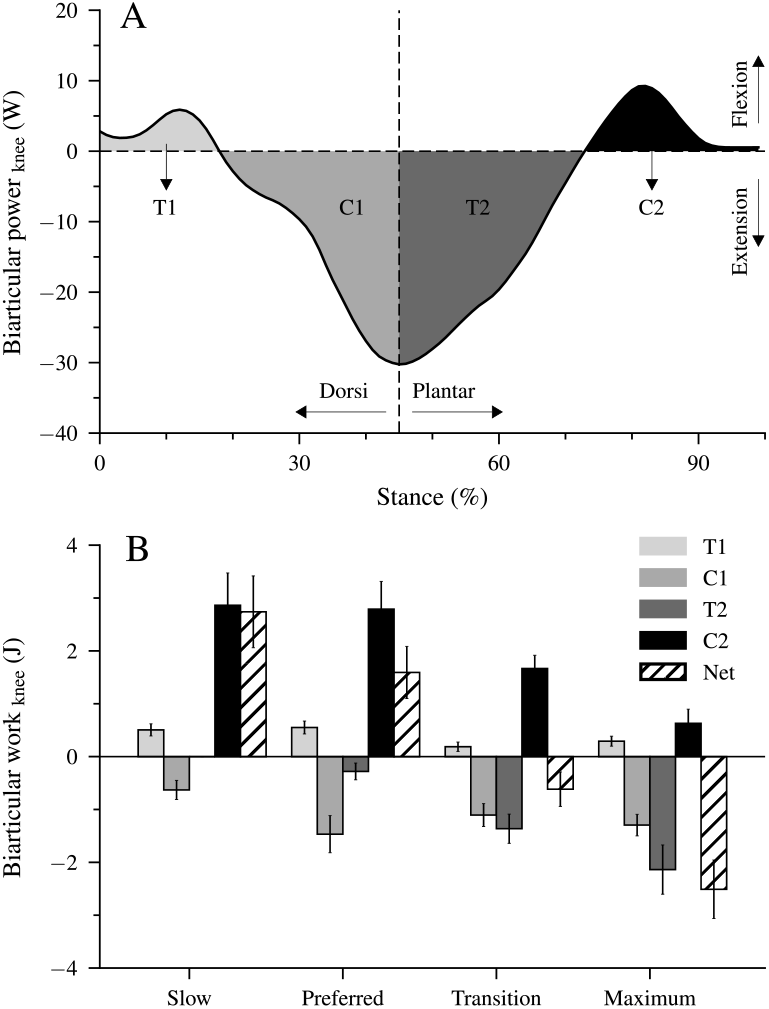
Mechanical power (A) and work (B) of the biarticular gastrocnemii muscles at the knee joint illustrating the transfer (T) and joint coupling mechanisms (C). T1: ankle-to-knee joint energy transfer, C1: energy absorption at the knee joint during dorsiflexion and knee extension, T2: knee-to-ankle joint energy transfer, C2: energy production at the knee joint during plantar flexion and knee flexion, Net: net work at the knee joint. The values are mean ± standard error.

## Discussion

Our results show that an increase in walking speed is associated with an increased net TSA-mechanical work, despite a decrease in the maximum AT-force in the transition and maximum compared to the preferred speed. We demonstrated that both the joint coupling and energy transfer mechanisms of the biarticular gastrocnemii muscles contributed to the speed-related enhanced net TSA-mechanical work by invariant net musculotendinous work. Furthermore, our findings show that in the two higher walking speeds, an earlier plantar flexion due to a rapid increase in the activation of the plantar flexor muscles was accompanied by a knee extension. The consequence was a knee-to-ankle joint energy transfer via the biarticular gastrocnemii muscles and earlier muscular energy production from the triceps surae muscles compared to the slow and preferred speeds. The continued elongation of the AT in the two higher speeds shows that a part of this energy transferred to the tendon despite the initiation of the plantar flexion. In line with our hypothesis, the results depict different mechanistic participation of the monoarticular soleus muscle and the biarticular gastrocnemii in the speed-related increased net TSA-mechanical work.

Similar to earlier studies (Neptune et al., 2008), we found a continuous decrease in stance and swing times and an increase in step length and cadence as a function of walking speed. The duty factor in all investigated speeds was >0.5, showing a double contact time at all speeds. The maximum AT-forces ranged between 1.616 and 2.174 kN and were very close to the measured values in comparable walking speeds reported by Finni et al. (Finni et al., 1998) using the optic fiber methodology. The maximum AT-force reduced by ~21% in the two higher speeds compared to the preferred one, despite a higher EMG activity, which confirms the predictions from musculoskeletal models at comparable walking speeds (Dominic James Farris & Gregory S Sawicki, 2012; Neptune & Sasaki, 2005). The increase of net TSA-work with speed can result from both an increased net mechanical work of the contractile elements of the triceps surae muscles and a modulation of the biarticular mechanisms. The most voluminous monoarticular soleus muscle can perform power and work only at the ankle joint. Based on the physiological cross-sectional area within the triceps surae muscles (Albracht et al., 2008), we assumed a 38% contribution of the gastrocnemii to the AT-force. This means that most of the speed-related increased net TSA-work is performed by the soleus muscle (i.e., 62%) or, more specifically, by the soleus contractile element, as the elastic element cannot produce positive net work. The unchanged net mechanical work of the gastrocnemii MTU indicates a similar net mechanical work of their contractile elements within the four investigated speeds. This implies that the contractile elements of the gastrocnemii muscles did not contribute to the speed-related increased net TSA-work.

The two gastrocnemii are biarticular muscles and, thus, are able to generate power and perform work in both ankle and knee joints. Due to their biarticularity, two additional mechanisms can affect the net TSA-work in addition to the work output of their contractile elements. The net mechanical work of the gastrocnemii muscles at the ankle joint showed a continuous increase from slow to the maximum speed of 4.3±0.8 J, demonstrating relevant participation of the biarticular mechanisms. This speed-related increase of the biarticular net work performed at the ankle joint without changes in the net work of the gastrocnemii MTU was due to a speed-related decrease in net mechanical work done at the knee joint. These findings provide first time evidence that the biarticular gastrocnemii muscles modulate their energy output within the two crossed joints towards an increased output at the ankle joint at higher walking speeds. Thus, the current study demonstrates that both the monoarticular soleus and the biarticular gastrocnemii contribute to the speed-related increased net mechanical work done at the ankle joint. However, the energetical processes and the mechanisms for this phenomenon differ between the two synergistic muscles. The soleus muscle enhanced the net mechanical work at the ankle joint by a speed-related increase of contractile work production (62% contribution), while the gastrocnemii muscles increased their work output at the ankle by an increased contribution of biarticular mechanisms (38%). The largest energy transfer from knee-to-ankle occurred at the maximum speed. More than three decades ago, van Ingen Schenau (Schenau, 1989) mentioned that the timing of the coupling between knee extension and plantar flexion could influence the effectivity of the knee-to-ankle energy transfer via the biarticular gastrocnemii muscles. The greater knee-to-ankle energy transfer in the maximum walking speed indicates a more effective coupling between knee extension and plantar flexion concerning energy production at the ankle joint compared to the transition and preferred speeds.

The earlier initiation of the plantar flexion in the two higher speeds (~47% of the stance phase) compared to the slow and preferred ones (~72% of the stance phase) was associated with a rapid increase in the EMG-activity of the Sol and GM muscles and with a knee-to-ankle energy transfer via the biarticular gastrocnemii muscles. This earlier initiation of the plantar flexion occurred at a low AT operating strain (Figure 6). At the beginning of the plantar flexion, the AT strain was 2.6±0.2% in the transition and 3.3±0.3% in the maximum walking speed, and both were clearly lower in comparison during slow 3.9±0.3% and preferred 4.1±0.3% walking speeds. The low AT-strain values in the two higher speeds indicate that the initiation of the plantar flexion and thus the production of mechanical work at the ankle joint occurred when the AT operates closer to or even within the toe region of the tendon force-elongation relationship, where the tendon may be elongated with rather low increments of muscle force. The energy transfer from the knee to the ankle via the biarticular gastrocnemii muscles and the rapid increase in triceps surae muscle activation at a low level of muscle forces (26±04% of the MVC) facilitated the triceps surae muscles force generation and storage of the elastic strain energy on the AT even during the plantar flexion. The simultaneous plantar flexion and AT elongation during the transition and maximum speeds show, at least for the monoarticular soleus muscle, a transfer of contractile energy production to the AT. Furthermore, the knee-to-ankle energy transfer via the gastrocnemii muscles indicates energy transfer from the proximal quadriceps muscles to the AT.

**Figure 6.**
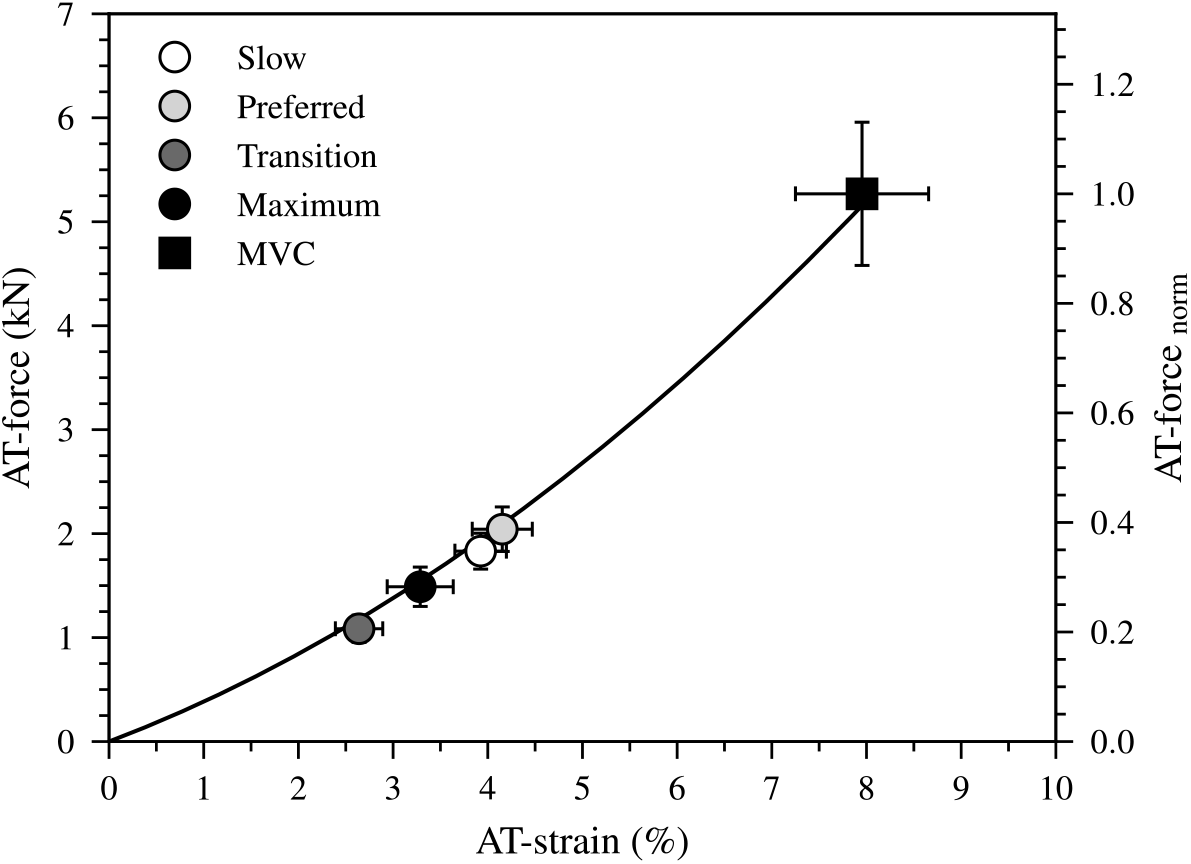
The force-strain relationship of the Achilles tendon (AT) during a maximum voluntary contraction (MVC). The markers show the operating AT-force and strain values at the beginning of the plantar flexion during walking at slow (0.7 m.s^−1^), preferred (1.4 m.s^−1^), transition (2 m.s^−1^) and maximum speed (2.6±0.3 m.s^−1^, Max) as means ± standard error.

The biarticular net work at the ankle joint increased as a function of walking speed and was greater than the net mechanical work of the gastrocnemii MTU, particularly at the two higher speeds. Also, the monoarticular muscle contractile energy production was greater in the transition and maximum walking speed compared to the preferred one. Although the contractile energy production of the monoarticular soleus and the participation of the biarticular mechanisms of the gastrocnemii muscles may increase the net mechanical work done at the ankle joint during the two higher speeds, and thus walking performance is metabolically costly. Contractile work during shortening increases the active muscle volume and thus the metabolic costs. The enormous increased EMG-activity of the investigated muscles in the transition and maximum walking speeds predict a higher active muscle volume. In the biarticular knee-to-ankle energy transfer mechanism, the more proximal quadriceps muscles are involved, and part of their contractile work can be transferred to the ankle joint (M. F. Bobbert et al., 1986; Gregoire et al., 1984). Due to the greater muscle volume of the quadriceps compared to the distal triceps surae muscles (Mersmann et al., 2014; Mersmann et al., 2015), they can produce more muscular power and work. However, due to their longer fibers, a unit of force generation is metabolically more expensive compared to the shorter triceps surae muscles (Biewener et al., 1998; Bohm et al., 2021). Therefore, the knee-to-ankle energy transfer at the two higher speeds might increase the metabolic costs of walking. Studies that measured the metabolic cost of walking show that the energy cost increased at speeds higher than the preferred and at speeds above 2.2 m.s^−1^, as our maximum speed (~2.6 m.s^−1^), it is even higher than during running (Minetti et al., 1994).

In conclusion, an increased net mechanical work performed at the ankle joint from the triceps surae muscles is needed to increase walking speed. The net mechanical work performed at the ankle joint can be affected by the contractile energy production of the triceps surae muscles and by the biarticular mechanisms of the gastrocnemii muscles. In the current study, we focused on the contribution of monoarticular and biarticular muscle mechanisms for the needed greater net TSA-work from slow to maximum walking speeds. Our findings show that both the contractile energy production in the monoarticular soleus muscle and the biarticular mechanisms of the gastrocnemii muscles contributed to the speed-related increased net mechanical work at the ankle joint. Although the contribution of the contractile work of the monoarticular soleus was greater, biarticular mechanisms of the gastrocnemii muscles accounted for a relevant part of the increased net ankle joint mechanical work with speed. An earlier plantar flexion initiated by a rapidly increased activation of the triceps surae muscles enhanced the net TSA-mechanical work 1.7 and 2.4-fold in the transition and maximum walking speeds, respectively, compared to the preferred one, despite a decrease in maximum AT-force.

## Methods

### Experimental Design

Fifteen healthy individuals (four female) with no record of musculoskeletal disorders (age 28±4 years, height 175.0±7.5 cm, body mass 75.0±9.5 kg) participated in the study. All participants submitted written informed consent to the experimental procedure that was approved by the ethics committee of the Humboldt-Universität zu Berlin (HU-KSBF-EK_2018_0005) and conducted in accordance with the Declaration of Helsinki. The participants were instructed to walk at the speeds of 0.7 m.s^−1^ (slow walking), 1.4 m.s^−1^ (preferred walking), 2.0 m.s^−1^ (transition walking) and their maximum walking speed capacity (2.6 ± 0.3 m.s^−1^) on a treadmill (Daum Electronic, Ergorun Premium8, Fürth, Germany). Individual maximum walking speed capacity was determined by gradually increasing the treadmill speed, starting from transition speed (i.e., 2.0 m.s^−1^), while they were instructed to maintain their walking pattern. When participants could not maintain the walking pattern and introduced a flight phase, the earlier selected speed was set as the maximum walking speed.

### Kinematics and electromyographic activity measurements

Six reflective markers (14 mm in diameter) were placed on anatomical landmarks, i.e., the tip of the second metatarsal, medial and lateral epicondyles of the femur, midpoint of a straight line between the greater trochanter and the lateral epicondyle of the femur and medial and lateral malleolus. One additional reflective foil marker (5 mm in diameter, flat surface) was placed on the insertion of the AT at the calcaneus (Figure 1). The insertion point of the AT was defined as the notch of the calcaneus bone, which was determined by moving a sound absorptive marker underneath an ultrasound probe in the sagittal plane until the shadow reflection (i.e., caused by sound absorptive marker) crossed the notch. Fourteen Vicon (Version 1.8.1, Vicon Motion Systems, Oxford, UK, 4 MX T20, 2 MX-T20-S, 6 MX F20, 2 MX F40, 250 Hz) cameras were used to capture the 3D trajectories of all markers in real-time. A fourth-order low pass, zero-phase shift Butterworth filter with a cut-off frequency of 12 Hz was applied to the raw marker trajectories. The participants performed a one-minute familiarization on the treadmill for each speed. The touchdown of the foot was determined as the heel marker’s instant minimal vertical position (Dingwell et al., 2001). The foot take-off was defined as the sign change in the second metatarsal marker velocity in the anterior-posterior direction (Alvim et al., 2015; Fellin et al., 2010). The ankle joint angle in the sagittal plane was calculated as the angle between the foot (i.e., line crossing the tip of the second metatarsal and calcaneus marker) and shank (i.e., line crossing ankle joint center and knee joint center). Ankle and knee joint centers were defined as the mean point between lateral and medial reflective markers on the malleoli and femur condyles. The knee joint angle in the sagittal plane was calculated as the angle between the femur axis (i.e., the line crossing the lateral knee and greater trochanter reflective marker) and the shank. The ankle and knee joint angles were calculated referencing a neutral, quiet stance position (the foot perpendicular to the tibia represents 0° ankle joint angle; the knee fully extended represents 0° knee joint angle). Positive values of ankle and knee joint angles constitute ankle plantar flexion and knee flexion.

A wireless system (Myon m 320RX, Myon AG, Baar, Switzerland) was used to measure the surface EMG activity of the TA, GM and Sol muscles during walking, operating at a sampling frequency of 1000 Hz. A fourth-order high-pass zero-phase shift Butterworth filter with a 50 Hz cut-off frequency, a full-wave rectification, and a low-pass zero-phase shift filter with a 20 Hz cut-off frequency was used to process the raw EMG signal. The resulting EMG signal was normalized to the highest processed EMG value acquired during a maximum voluntary isometric contraction (MVC).

### Measurement of AT-length and quantification of AT-force and strain energy

The origin of the AT was determined as the most distal junction between the AT and the medial head of the gastrocnemius muscle (GM-MTJ), obtained by transverse and sagittal ultrasound scans (Figure 1). A 60 mm T-shape ultrasound probe (Aloka UST-5713T, Hitachi Prosound, alpha 7, Japan) with a sampling frequency of 146 Hz was fixed above the GM-MTJ and was tightened firmly with a flexible plastic cast. An ultrasound gel pad was placed between the ultrasound probe and the skin to compensate for the unevenness of the skin surface. The ultrasound and motion-capture systems were synchronized using a 5-volt manual trigger through the ultrasound electrocardiograph channel and motion-capture analog input channel.

A semi-automatic image-based tracking algorithm was implemented in a self-developed user interface (MATLAB, version 9.6. Natick, Massachusetts: The MathWorks Inc) to track the position of the GM-MTJ from the stack of ultrasound images. The details of the developed algorithm and its validity compared with manual tracking during human walking and running (i.e., r^2^=0.97) were provided earlier (Kharazi et al., 2021). The skin surface was detected using a double threshold-to-intensity gradient provided by the Canny edge detection function (Canny, 1987). A third-order polynomial curve was then fitted to the detected points on the skin, and the tracked coordinates of the GM-MTJ were perpendicularly (i.e., shortest distance) projected to the fitted curve.

For the transformation of the GM-MTJ to the global coordinate system, the four corners of the ultrasound probe’s protective plastic rubber were digitized in 3D space using a custom 3D-printed calibration tool in a separate session. A coordinate system was defined on the left-center side of the plastic rubber. The gap between the origin of the specified coordinate system and the first piezoelectric underneath was determined by subtracting the ultrasound rubber plastic width from the ultrasound image width. A custom 3D-printed tripod was mounted on the ultrasound probe, and a coordinate system was defined accordingly. The projected positions of the GM-MTJ on the skin surface were then transferred to the global motion capture’s coordinate system (for details, see (Kharazi et al., 2021). Depending on the location of the GM-MTJ, reflective foil markers with 20 mm intervals were placed on the skin from the calcaneus bone on the path of the AT to the last possible position below the cast (Figure 1). The curved length of the AT was determined as the sum of vectors in every two consecutive points using the 3D coordinates of the foil markers and the GM-MTJ projected to the skin. The strain of the AT was calculated by dividing the measured AT elongation by the AT length measured at a relaxed state in 20° plantar flexion, where AT slackness has been reported earlier (De Monte et al., 2006).

Potential displacements of the skin to the bone underneath the calcaneus marker that defines the AT insertion can also introduce artifacts in the AT-length measurement (Lichtwark & Wilson, 2005). In a separate experiment, kinematic and ultrasound measurements were used to measure the displacement of the reflective calcaneus marker above the defined insertion point (notch at tuber calcanei) to account for the potential movements of the skin relative to the calcaneus bone (Kharazi et al., 2021). For this purpose, a sound absorptive marker was placed between the ultrasound probe and skin close to the bony insertion and the dynamometer (Biodex Medical, Syst.3, Shirley, NY) passively rotated the ankle joint. The displacement of the calcaneus bone relative to the skin as a function of the heel angle (angle between the line from the calcaneus marker to ankle joint center and the line from the ankle joint center to knee joint center) was determined and used as a model to correct the AT-length during walking as a function of the heel angle. The potential error due to the skin-bone artifact was 0.92±0.57 mm (mean ± standard deviation) on the AT-length and 0.45% on AT-strain (Kharazi et al., 2021).

In an additional session, an individual force-elongation relationship of the AT was determined. For the measurement of the plantar flexion moment, five ramp MVC (~ 5 seconds gradual increase in a moment) plantar flexions were performed at a joint angle position of 0° and fully extended knee on a dynamometer. The ankle and dynamometer axes of rotation misalignments during the contractions, and gravitational and passive moments were considered by employing an inverse dynamics approach (Arampatzis et al., 2005). Further, we considered the effect of co-activation of the antagonistic muscle (i.e., tibialis anterior) on the resultant joint moment during the MVCs using an established EMG-based approach (Mademli et al., 2004). The ankle joint moment was then divided by the AT lever-arm to calculate the tendon force. For this purpose, the tendon lever-arm was calculated using the tendon-excursion method and adjusted for the changes in the alignment of the AT during the contractions using the data provided by Maganaris et al. (Maganaris et al., 1998). A 10-centimeter linear ultrasound probe (My lab60, Esaote, Genova, Italy, 25 Hz) was fixed above the GM-MTJ to measure the corresponding elongation of the AT during the five MVCs. The GM-MTJ displacement resulting from ankle joint rotations (i.e., during MVC) was considered by tracking its displacement during a passive rotation (at 5°/s) of the ankle joint (Arampatzis et al., 2008). To achieve excellent reliability, the tendon force-elongation relationship of each participant was averaged from five MVCs (Schulze et al., 2012). The individual force-elongation relationship of the AT was obtained by fitting a quadratic function (Eq 1), which was then used to assess the AT force during walking.

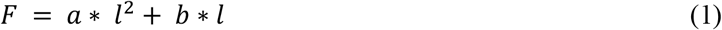

Where F is the AT-force, *a* and *b* are the coefficients of the quadratic function, and *l* is the elongation of AT during the MVCs.

Using Eq 1 and the measured AT-elongation during the walking trials, we calculated the AT-force and then AT elastic strain energy as the integral of the AT-force over AT-elongation (Eq 2) during the stance phase.

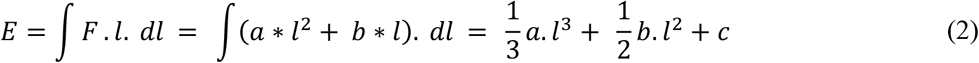

Where the *E* and *F* are the AT strain energy and force. *l* is the instantaneous elongation of AT during stance, and *c* is the constant of the integral. The AT power was calculated as the first time-derivative of the AT elastic strain energy.

### Triceps surae mechanical power and work performed at the ankle and knee joints

The ankle joint moment resulting from the triceps surae muscles (TSA-moment) during walking was calculated as the product of the AT force and the AT lever arm. The instant AT lever-arm was measured as the perpendicular distance between the AT line of action and the marker-based ankle joint center during the walking trials (Figure 1). The line of action of the AT was assumed as the midpoint of the AT. The distance between the skin surface and the midpoint of the AT was assessed using sagittal plane ultrasound scans and subtracted from the kinematic-based AT lever arm. The ultrasound probe was placed above the AT on the level of the malleoli while participants were seated in a relaxed prone position with their ankle angle set at 0°. The depth between the skin surface and the midpoint of the AT among all individuals was, on average, 5.0±0.3 mm. The mechanical power performed at the ankle joint from the triceps surae muscles (TSA-power) was calculated as the product of TSA-moment and ankle angular velocity. The mechanical work done from the triceps surae muscles (TSA-work) at the ankle joint was calculated as the integral of TSA-power over time.

The GM and GL, as biarticular muscles, can generate or absorb power and energy in both the ankle and knee joints. This means they can redistribute the mechanical power and work performed by their MTUs over the two joints and transfer power and energy from knee-to-ankle and vice versa. We investigated the mechanical power and work performed at the ankle and knee joints by the gastrocnemius medialis and gastrocnemius lateralis muscles in order to examine their contribution to ankle and knee mechanical power and work and the energy transfer between the two joints during walking. Mechanical power at the ankle/knee joints was calculated as the product of the ankle/knee joint moment generated from the two muscles and ankle/knee joint angular velocity. The generated moment at the ankle/knee joint by the two muscles was calculated as the product of the gastrocnemius medialis and gastrocnemius lateralis muscle force and their instantaneous moment arms. The gastrocnemius medialis and gastrocnemius lateralis muscle forces have been assessed as a fraction of the AT-force by using the relative physiological cross-sectional area of the two muscles with respect to the triceps surae muscle (i.e., 26 % for the gastrocnemius medialis and 12% for the gastrocnemius lateralis) as reported by Albracht et al. (Albracht et al., 2008), assuming the force contribution of each muscle is proportional to its physiological cross-sectional area. The moment arm of the gastrocnemius medialis and gastrocnemius lateralis at the knee joint were extracted as a function of the knee joint angle using the values reported by Buford et al. (Buford et al., 1997). The mechanical work of the gastrocnemii muscles performed at the ankle and knee joint was then calculated as the integral of the ankle and knee joint power over time. The sum of the mechanical power/work performed at the ankle and knee joints by the biarticular gastrocnemii equals the mechanical power/work performed by the gastrocnemii MTU (Prilutsky et al., 1996).

### Statistics

A linear mixed model was used to test for the main effect of walking speed on all investigated outcomes (i.e., temporal and spatial gait parameters, kinematics, EMG, strain, moments, force, and mechanical work). The significance level was set to *α* = 0.05 and all values are reported as mean ± standard errors. A pairwise Tukey-test was performed as a post hoc analysis in case of a significant main effect of speed, and Benjamini-Hochberg corrected *p*-values will be reported. The statistical analyses were conducted using R v4.0.1 (R foundation for statistical computing, Vienna, Austria. Packages), where the “nlme” package was used for the linear mixed model and the “emmeans” package for post hoc testing.

## Acknowledgements

We acknowledge the German Research Foundation (DFG) support and the Open Access Publication Fund of the Humboldt-Universität zu Berlin.

## Author contributions

Mohamadreza Kharazi participated in designing the methods, carried out the experiments and data analysis and drafted the manuscript; Sebastian Bohm coordinated and participated in designing the study, carrying out the experiment, reviewed and edited the manuscript; Chris Theodorakis participated in the experiment and data acquisition, reviewed and edited the manuscript; Falk Mersmann participated in language proofing of the manuscript, reviewed and edited the manuscript and Adamantios Arampatzis conceived, designed and coordinated the study and drafted the manuscript. All authors gave final approval for publication.

## Ethics

Human subjects: The ethics committee of the Humboldt-Universität zu Berlin approved the study and the participants gave written informed consent in accordance with the Declaration of Helsinki.

